# Community metabolic modeling of host-microbiota interactions through multi-objective optimization

**DOI:** 10.1101/2023.09.15.557910

**Authors:** Anna Lambert, Marko Budinich, Maxime Mahé, Samuel Chaffron, Damien Eveillard

## Abstract

The human gut microbiota comprises various microorganisms engaged in intricate interactions among themselves and with the host, affecting its health. While advancements in omics technologies have led to the inference of clear associations between microbiome composition and health conditions, we usually lack a causal and mechanistic understanding of these associations. For modeling mechanisms driving the interactions, we simulated the organism’s metabolism using *in silico* Genome-Scale Metabolic Models (GEMs). We used multi-objective optimization to predict and explain metabolic interactions among gut microbes and an intestinal epithelial cell. We developed a score integrating model simulation results to predict the type (competition, neutralism, mutualism) and quantify the interaction between several organisms. This framework uncovered a potential cross-feeding for choline, explaining the predicted mutualism between *Lactobacillus rhamnosus* GG and the epithelial cell. Finally, we analyzed a five-organism ecosystem, revealing that a minimal microbiota can favor the epithelial cell’s maintenance.

## Introduction

The human gut microbiota is a complex ecosystem that significantly impacts host health and disease across the life course (1). Therefore, clarifying its role and associated mechanisms in shaping health is an essential fundamental research goal. Environmental omics enable a better understanding of this ecosystem by studying its composition and variation associated with host phenotypes (2). This has led to identifying bacteria promoting health (e.g., *Faecalibacterium Prausnitzii* is depleted in patients with Inflammatory Bowel disease (3)) or disease (e.g., Class *Betaproteobacteria* is enriched in diabetic individuals (4)). However, correlation does not imply causality, and each strain’s mode of action is yet to be understood. Disentangling the impact of bacteria within a diverse ecosystem structured by heterogeneous environmental and host factors is challenging.

Gut microbiota metabolism is central to host physiology as its participation in digestion modulates nutrient availability to the host (5, 6). Moreover, the cross-talk among bacteria shapes the ecosystem (7), consequently modulating its function (8). To investigate mechanistic metabolic interactions between organisms, computational models are useful tools (9, 10). Researchers can reconstruct metabolic networks based on an organism’s complete genome, biochemical databases, and literature knowledge. Leveraging stoichiometry and thermodynamics information, this network can be formalized into a Genome-Scale Metabolic Model (GEM), facilitating the simulation and examination of its metabolic phenotype (11). Indeed, GEMs predict the metabolic phenotype in a given condition and explain this prediction by revealing important nutrients and activated metabolic pathways (12). Many GEMs are now available (13, 14), and automated reconstruction processes for microorganisms have emerged (15–17), paving the way to analyze various potentially unexplored bacterial metabolic mechanisms. Simulating the metabolism of individual organisms is a well-established and extensively employed approach for expanding fundamental knowledge. For instance, it aids in closing gaps in metabolic knowledge by identifying differences between predictions and experimentations (11). Additionally, this approach contributes to enhancing industrial capacity. An example of this is genetic manipulation, which boosts the production of molecules of interest (18). To orient the model toward biologically relevant metabolic behavior, its biomass production, representing its growth, is maximized (19), emphasizing phenotypes where the organism undergoes replication. For a community, this means maximizing the overall biomass produced by all organisms in the ecosystem. While interesting, this method favors the organism with the better yield, possibly outcompeting the others. We expect a more nuanced compromise in biological systems, driven by metabolic trade-offs and resource competition (20). To avoid this bias, the ecosystem biomass is generally weighted based on the relative abundance of each species in a specific condition (e.g., the Microbiome Modeling Toolbox (21)), aligning their growth rate to their effective presence in the ecosystem. To go further, Diener et al. introduced MICOM (22), a framework inferring the growth rate from relative abundance data, refining the weights of the ecosystem biomass. This method calibrates the model to fit observed species proportions inferred from omics data. Parametrizing the model to better simulate a known condition allows for satisfactory predictions with simpler models, but the dependence on environmental data is troublesome. Here, we aimed to achieve non-parametric modeling, striving for accurate predictions through refined model analysis rather than from model fitting.

We used multi-objective linear programming to predict and explore potential interactions between several organisms without restricting ourselves to a unique configuration. This approach allows for the optimization of independent objectives, revealing trade-offs between each organism’s biomass as a Pareto front. This is usually applied in bi-objective modeling of microbial ecosystems to infer the type of interaction between bacteria of interest, which is made accessible through the Metabolic Modeling Toolbox (21).

Here, we deployed a multi-objective modeling framework to gain further insights into the interaction between gut microbes and small intestinal epithelial cells. We inferred an interaction score predicting the type and level of interaction among organisms within an ecosystem based on the analysis of the Pareto front. This analysis highlighted known probiotics when applied to 331 bacterium-enterocyte ecosystems. Furthermore, we delved into the mechanisms underlying the mutual interaction between *Lactobacillus rhamnosus* GG (LGG) and the enterocyte, uncovering a potential cross-feeding relationship involving choline. Finally, we integrated four bacterial models with the enterocyte model to explore the intricate metabolic interplays within a more complex ecosystem. In this context, our findings reveal that the presence of gut bacteria significantly influences and supports the enterocyte’s objective (i.e., maintenance of the cell without replication).

## Results

### Scoring metabolic interaction of bacterium-enterocyte ecosystem models using bi-objective optimization

To explore metabolic interactions between the human gut and the microbiota, and quantify them as a score, we built ecosystem models defined as the integration of GEMs of small intestinal epithelial cells (i.e., enterocyte) and one or several bacteria through a pool compartment (Figure 1A). Classically, metabolic modeling allows the maximization of a single objective: the growth of the bacteria or the maintenance of the enterocyte (Figure 1B). The plurality of objectives must be addressed when joining several organisms in an ecosystem. To simulate ecosystem behaviors, we consider that each organism independently maximizes its objective, shaping the competition and cooperation for available nutrients (23). It results in a problem one can solve using multi-objective linear programming. The set of all optimal solutions for the ecosystem is a Pareto front. It reflects the trade-offs between all involved biological objectives. Herein, we advocate that the shape of the Pareto front describes the nature and level of metabolic interaction between the organisms and can be summarized as a score. As a first approximation, we worked with simple ecosystems consisting of a single bacterium and the enterocyte. In this context, the Pareto front comprises two dimensions: one for each objective. Maximal growths of each organism alone are used to normalize each axis as they standardize growths under interaction regimes (i.e., studying the interaction between organisms without an influence of the original growth). This transformation is essential to make our interaction score comparable and interpretable between ecosystems built using different bacteria. When both objectives negatively or positively impact the other, both organisms are in competition or mutualism, respectively. When neither objective value is affected by the other, it is considered neutralism (see Figure 1C to illustrate three representative Pareto front shapes). We propose an Ecosystem Interaction Score (S) based on the normalized Pareto front’s Area Under the Curve (AUC). The AUC of the “non-interaction front,” defined by the neutral interaction, is subtracted from the AUC of the normalized Pareto front. It results in a positive score for mutualism, a null score for neutrality, and a negative score for competition (see Methods for details).

**Fig. 1.**
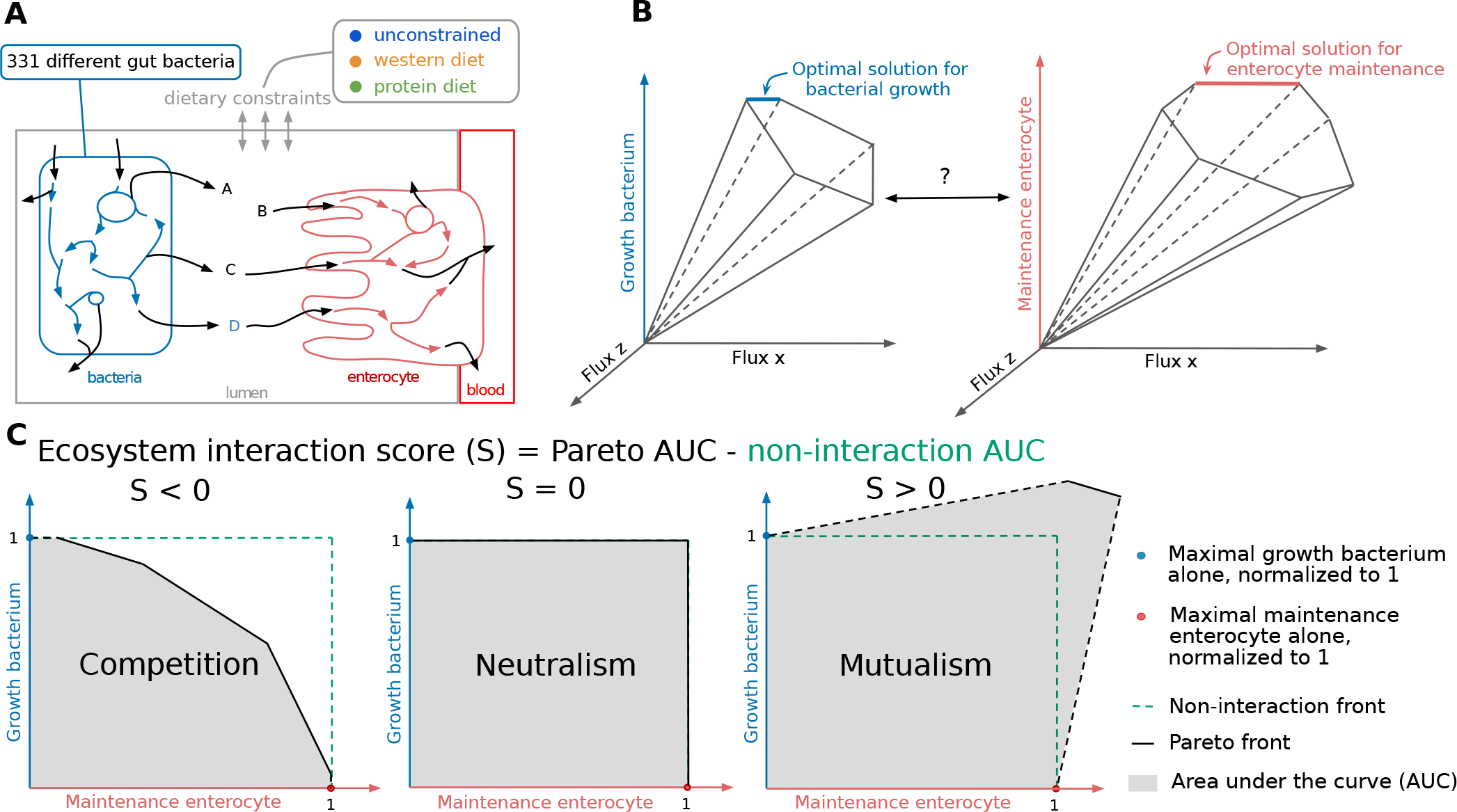
Schematic rationale of the Genome-Scale modeling and its application in the metabolic interaction between two systems within an ecosystem. **(A)** Representation of an ecosystem model built from a bacterium (blue) and the enterocyte’s (red) GEMs joined together through a lumen compartment (grey). Arrows represent metabolic reactions. Black arrows are transport reactions, enabling the transit of metabolites between compartments. Three different diets (unconstrained, western diet (WD) and protein diet (PD)) were imposed on the ecosystem by defining corresponding dietary constraints (see Methods). **(B)** Solving a GEM implies identifying its solution space and optimal solutions when maximizing its objective function, often resumed by its growth rate. The solution space of a bacterium (left) described here in three dimensions illustrates how the optimal solutions for the bacterium were identified by maximizing its growth. Similarly, the optimal solutions for the enterocyte are identified by maximizing its cell maintenance (cellular membrane maintenance, proteins and energy production, no replication). Both objectives must be accounted for in an ecosystem model, which is possible using multi-objective optimization. **(C)** Schematic representation of three different Pareto fronts and how it can be used to define the ecosystem interaction score (S). The shape and area of the bi-objective solution space can be used to define competition (left panel), neutralism (center panel, defining the non-interaction front), and mutualism (right panel). Dotted lines link the added “original growth points” and the strict Pareto front.

### Interaction score for 331 gut bacterium-enterocyte ecosystems under three different dietary conditions

331 strain-level gut bacteria models from the EMBL GEMs (16) were selected as they were described as gut microbes based on the Virtual Metabolic Human (VMH) Database (14). Their pairwise interaction score with the enterocyte was computed under three nutritional conditions (Figure 1A, Table S1 for complete results). The first condition, an unconstrained diet, represents a synthetic environment where all modeled nutrients are unlimited in the lumen. The other two conditions, the Western Diet (WD) and the protein diet (PD), are biologically representative diets. The WD is high in fat and simple sugars but low in fibers, while the PD is rich in protein but poorer in fats and simple sugars than the WD (24) (see Methods). An increased number of non-dominated points on the Pareto front reveals more factors affecting the interaction, characterizing its complexity.

Overall, ecosystems had an interaction score close to zero under the unconstrained diet but exhibited negative and positive scores under the WD and PD (Figure 2A). This greater predicted interaction observed in constrained media was expected, illustrating either the mutual reliance on a common metabolite leading to competition or the adaptation through cross-feeding, usually favored in nutrient-limited environments (25).

**Fig. 2.**
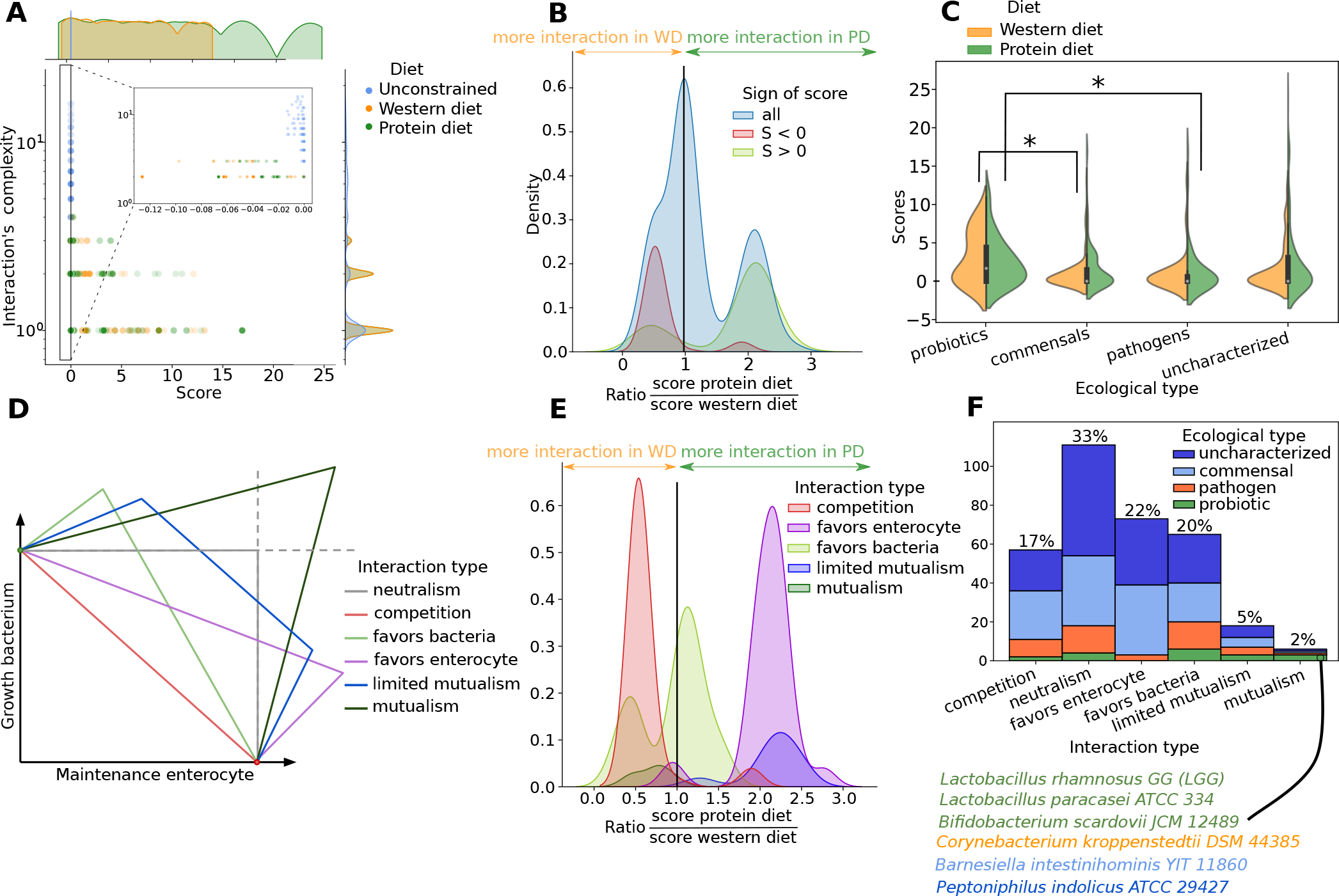
Prediction of metabolic interaction scores and types for 331 enterocyte-bacterium ecosystems. **(A)** Interaction score (S) 2D distribution and density curve among 331 ecosystems subject to three diets (unconstrained, WD, PD), with a zoom on negative values. The number of points constituting the Pareto front defines the interaction complexity. **(B)** Evaluation of the effect of the diet on the score through a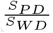 distribution colored by the sign of the score. **(C)** Assessment of the predictive potential of S on the ecological type (probiotics n = 18, pathogens n = 46, commensals n = 123, uncharacterized n = 144) colored by diet (WD or PD). A Mann-Whitney U-test was used to determine statistical significance. **(D)** Illustration of the interaction types inferred from the Pareto shapes (see Methods). **(E)** Evaluation of the impact of diet on the score in interaction types using 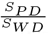 ratio distribution. **(F)** Proportions of each Pareto inferred interaction type and exploration of how ecological types place themselves on the interaction types. Specification of bacterial strain predicted as mutualist with the enterocyte.

The sensitivity of the interaction score to diets is demonstrated by observed differences in scores between PD and WD for a given ecosystem. Furthermore, ecosystems with positive scores generally had an increased score under PD compared to WD, while those with negative scores showed a decrease in score under WD compared to PD (Figure 2B). In other words, the PD appeared to enhance positive interactions between bacteria and the enterocyte, while the WD is predicted to favor negative interactions.

### Further characterization of the Pareto front into “interaction types”

The shapes of the Pareto fronts are more diverse than depicted in Figure 1C. They can inform us about the nature of biotic interactions. To leverage this information and enrich the metabolic interaction score, we defined discrete categories as “interaction type” (Figure 2D and see Methods). Indeed, a score can be positive but only favoring one organism, as represented by the interaction types “Favors bacteria” and “Favors enterocyte.” In some cases, the presence of the other can favor both organisms, but this advantage disappears as the other organism gets to maximize its objective, as observed in “Limited mutualism.” “Competition,” “Neutralism,” and “Mutualism” interaction types correspond to the shapes described in Figure 1C.

To determine how S varied based on diet in each interaction type, we distributed the ratio calculated previously 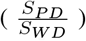 by interaction types (Figure 2E). Most positive interactions, specifically those favoring the enterocyte (favors enterocyte, limited mutualism), were increased in PD. However, mutualistic interactions were more important in WD. Interestingly, the increase of S in WD for the mutualistic ecosystem is not explained by reaching a higher objective value but because the interaction partially compensates for the lower objective value of each organism in WD compared to PD (Table S2). More than a beneficial effect of WD, this illustrates how limited resources can increase interactions (25). While WD enhanced some interactions favoring bacteria, it mostly favored competitive interactions.

### Integration of metabolic interaction score and type to predict “Host-beneficial” microbes

The VMH database (14) provided the ecological type of the 331 bacterial GEMs. Microorganisms were categorized into three ecological types: probiotic, pathogen, commensal, and uncharacterized. Probiotics had a significantly higher score compared to pathogens in PD (Mann-Whitney, p = 0.033) and commensals in WD (Mann-Whitney, p = 0.040) (Figure 2C). Overall, we observed a tendency for higher scores in known probiotics (Figure 2C). This result raises interest in some uncharacterized bacteria displaying a high interaction score with the enterocyte as Cetobacterium somerae ATCC BAA-474 (S_*P D*_ = 23.64, S_*W D*_ = 12.10), Klebsiella aerogenes KCTC 2190 (S_*P D*_ = 16.92, S_*W D*_ = 8.70) or Morganella morganii subsp morganii KT (S_*P D*_ = 11.42, S_*W D*_ = 5.80).

Next, we explored the ecological type distribution among interaction types (Figure 2F). A third of the simulated bacteria (33%), encompassing all ecological types, were predicted to have a neutral interaction with the enterocyte. Many bacteria favored the enterocyte (22%), largely dominated by commensals and uncharacterized bacteria. Ecosystems where the bacteria’s growth was favored (20%) contained more pathogens and probiotics than the ones favoring the enterocyte. 17% of the bacteria were engaged in a competitive interaction with the host. Limited and high mutualisms were identified as the least common interaction types, accounting for only 5% and 2% of the analyzed bacteria, respectively. Probiotics corresponded to half of the mutualistic bacteria (N=6) with two lactic acid bacteria and a short-chain fatty acid producer (*Lactobacillus rhamnosus* GG, *Lactobacillus paracasei* ATCC 334, and *Bifidobacterium scardovii* JCM 12489). Additionally, unexpected mutualistic interactions were observed for *Corynobacterium kroppenstedtii* DSM 44385 (categorized as a pathogen), *Barnesiella intestinihominis* YIT 11860 (classified as a commensal), and *Peptoniphilus indolicus* ATCC 29427 (tagged as uncharacterized) with the enterocyte.

### Metabolic exchanges driving mutualism between the enterocyte and *Lactobacillus rhamnosus* GG reveal a potential cross-feeding of choline

*Lactobacillus rhamnosus* GG (LGG) is a very well-studied probiotic bacteria. Among other properties, it shows good adherence to the intestinal epithelial layer and supports the survival of intestinal epithelial cells (26). Our computational analysis predicts LGG to engage in a mutualistic relationship with the host, as reflected by its interaction scores of 8.68 in WD and 6.60 in PD. Their Pareto front formed a spike from which the peak allowed the highest biomass production for both organisms and, consequently, for the ecosystem (See Table S2). To unravel the underlying mechanisms driving this mutualistic interaction by identifying essential reactions to reach this optimum, we sampled 100,000 solutions on the Pareto front in WD. We focused on exchange reactions that exhibited a strong correlation (corr ≥ 0.95, Spearman) with the overall biomass production of the ecosystem (i.e., summed objective values, here, the peak of the Pareto front), as depicted in Figure 3A.

**Fig. 3.**
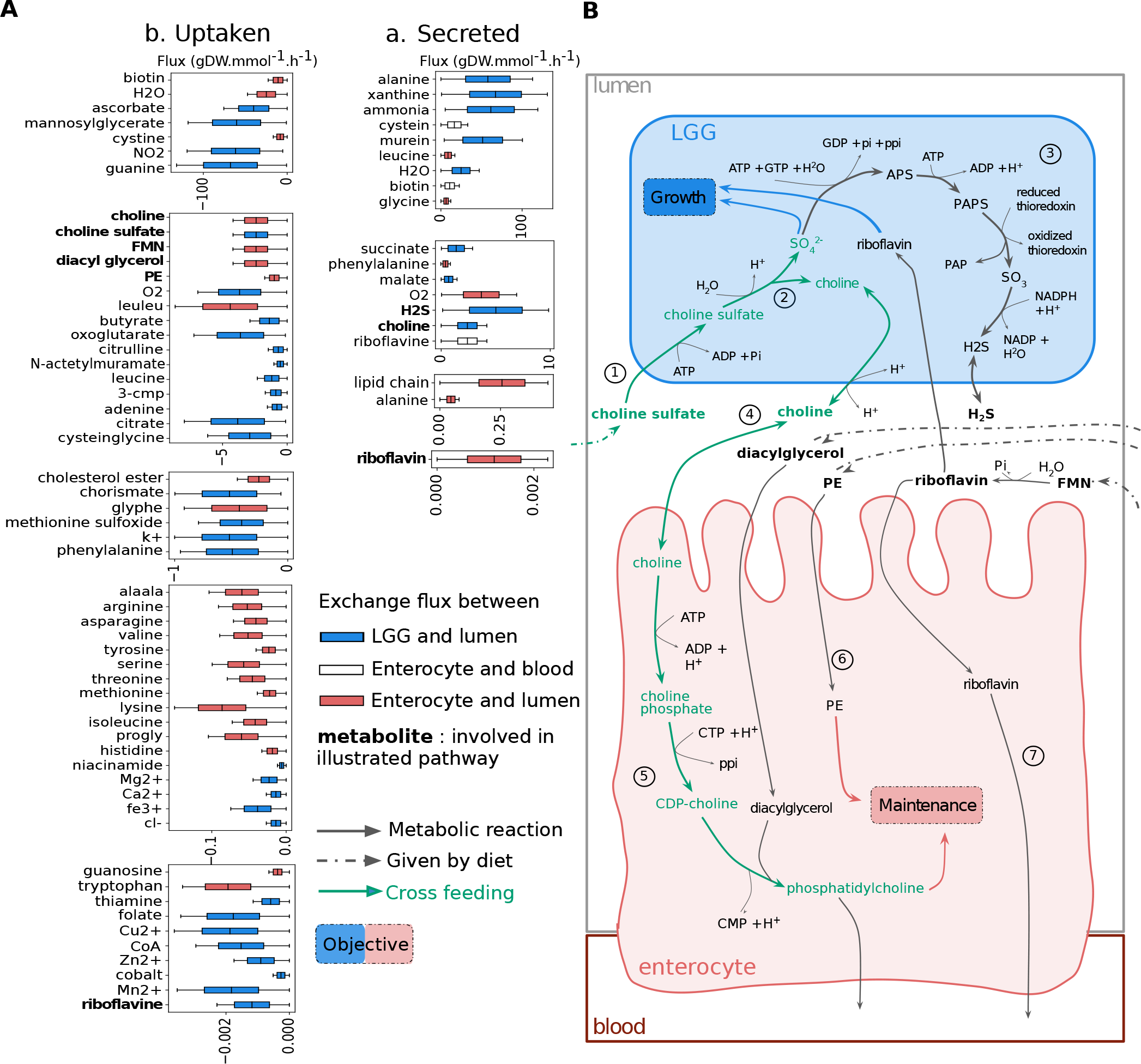
Exploring the Pareto front of the LGG-enterocyte interaction reveals a possible cross-feeding involving choline. **(A)** We Sampled the Pareto front of the interaction between LGG (blue) and the enterocyte (red). We inferred the correlations between transport reaction and ecosystem biomass to identify metabolic cross-feedings driving the interaction. Metabolites for which the exchanges are correlated with the ecosystem biomass, therefore driving the interaction, are described in boxplots by the distribution of their possible fluxes among the sampling. A metabolite is categorized as uptaken (or secreted) when its absorption (or excretion) by the relevant organism is associated with ecosystem biomass (see Methods). The color of the boxplot refers to the relevant organism (blue for LGG, red for the enterocyte). White boxplots refer to metabolites transported into the blood from the enterocyte. Metabolites involved in the metabolism depicted in (B) are in bold. **(B)** Schematic representation of the predicted cross-feeding of choline. The arrows represent metabolic reactions. Dashed arrows represent the exchange reactions (i.e., the consumption of metabolites from the diet). Blue and red arrows illustrate the use of a metabolite for the objective of the LGG and the host, respectively. The metabolic pathway colored in green is the predicted cross-feeding of choline. All reactions pictured in this figure are obligatory (See Methods) in the most mutualistic solution of the Pareto front.

The exchange reactions identified through this analysis enabled both organisms to reach higher objective values when interacting, helping to predict valuable nutrients and cross-feeding pathways. The intricate metabolic interplay within the ecosystem involves the shared utilization of various metabolites, contributing to the overall ecosystem functioning. For the ecosystem to reach optimal biomass, we predicted the uptake of many amino acids and dipeptides to be essential, mainly for the enterocyte. However, the digestion of leucine-leucine and glycine-phenylalanine dipeptides by the enterocyte’s enzymes provided the bacteria with leucine and phenylalanine. Oligoelements and vitamins were more important for LGG, but the enterocyte uptaked biotin and riboflavin (7) and released them in the blood. The secretion of choline by LGG was found to be strongly correlated with its uptake by the enterocyte (corr = 0.96, Figure S2), and both these transports are associated with a higher biomass for the ecosystem. Specifically, as illustrated in Figure 3B, our model predicted that LGG uptakes choline sulfate (1), a compound found in human food such as plants, algae, and numerous fungi such as *Aspergillus oryzae*, a ferment used for sake, miso, and soy sauce (27, 28). LGG hydrolyzes choline sulfate (2) in sulfate ions 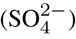, which are useful for LGG’s growth, and choline, secreted in the intestinal lumen. LGG reduces the excess sulfate to H_2_S (3) and excretes it in the lumen. The choline, now available in the lumen, is uptaken by the enterocyte (4). In the enterocyte, the absorbed choline transforms into CDP-choline (5) which, along with diacylglycerol, further converts to phosphatidylcholine. Phosphatidylcholine is the principal component of cell membranes and participates in lipid metabolism (29), therefore regarded as an essential brick for cell maintenance.

Choline is an essential nutrient and source of methyl. However, less than half of the tested populations (adult men and women) reached the recommended intake (30). Therefore, it is consistent with the literature to expect its supplementation, through cross-feeding with LGG, to favor the enterocyte’s maintenance. Phosphatidylethanolamine (PE), another abundant phospholipid (29), was available in excess in the environment when the enterocyte was simulated on its own. However, since phosphatidylcholine was limited in availability, the enterocyte’s maintenance and PE intake were constrained. In the community with LGG, as the choline availability increased, the enterocyte also exhibited an elevated absorption of PE (6), conjointly leading to improved maintenance for the enterocyte.

### Modeled minimal gut microbiome metabolism’s greatly favors the enterocyte

An ecosystem is formed of more than two organisms, and to move toward modeling realistic gut-microbiota interactions, more organisms must be included in the modeled community. The bacterial strains from Shetty et al.’s minimal microbiome (31) were modeled using CarveMe (16), and four were selected because of their distinct metabolic potential and predicted phenotypes: *Akkermansia muciniphila* ATCC BAA-835, *Bacteroides xylanisolvens* HMP 2_1_22, *Faecalibacterium prausnitzii* A2-165 and *Ruminococcus bromii* ATCC 27255. It should be noted that while four bacteria were chosen for this analysis, it is computationally applicable to up to 10 bacteria and the enterocyte (Figure S3). They were integrated with the enterocyte in an ecosystem model, which was analyzed using Multi-objective linear programming. Here, as the Pareto front is in five-dimensional space, its description was reduced to its extreme points for analytical purposes, constituting the extreme solutions of the ecosystem.

Solutions from the Pareto front represent extreme community phenotypes where all available nutrient usage is optimized. To envision a more realistic (i.e., suboptimal) set of community phenotypes, three thousands additional solutions were randomly sampled within the entire solution space (i.e., solutions within the volume embedded by the Pareto front, such as the grey surface in Figure 1 when applied in 2 dimensions). Combining optimal and random solutions results in an inclusive set of potential phenotypes for the ecosystem. Here, solutions are described by a 5-dimensional vector, each value being an organism’s objective. To visualize how each organism’s objective impacts the rest of the ecosystem, we performed a Principal Component Analysis (PCA) on the objective values (Figure 4A). The PCA revealed that PC1, which explains 30% of the variance, strongly aligns with an increase in ecosystem biomass and is coherently bordered by extreme solutions. Notably, all organisms contributed to the increase in ecosystem biomass. PC2 (21% of explained variance) refined this interpretation and distinguished distinct groups within the ecosystem: *Bacteroides xylanisolvens*, and the enterocyte exhibited a similar trend, while *Akkermansia muciniphila* showed an opposing direction. *Ruminococcus bromii* and *Faecalibacterium prausnitzii* fell between these groups.

**Fig. 4.**
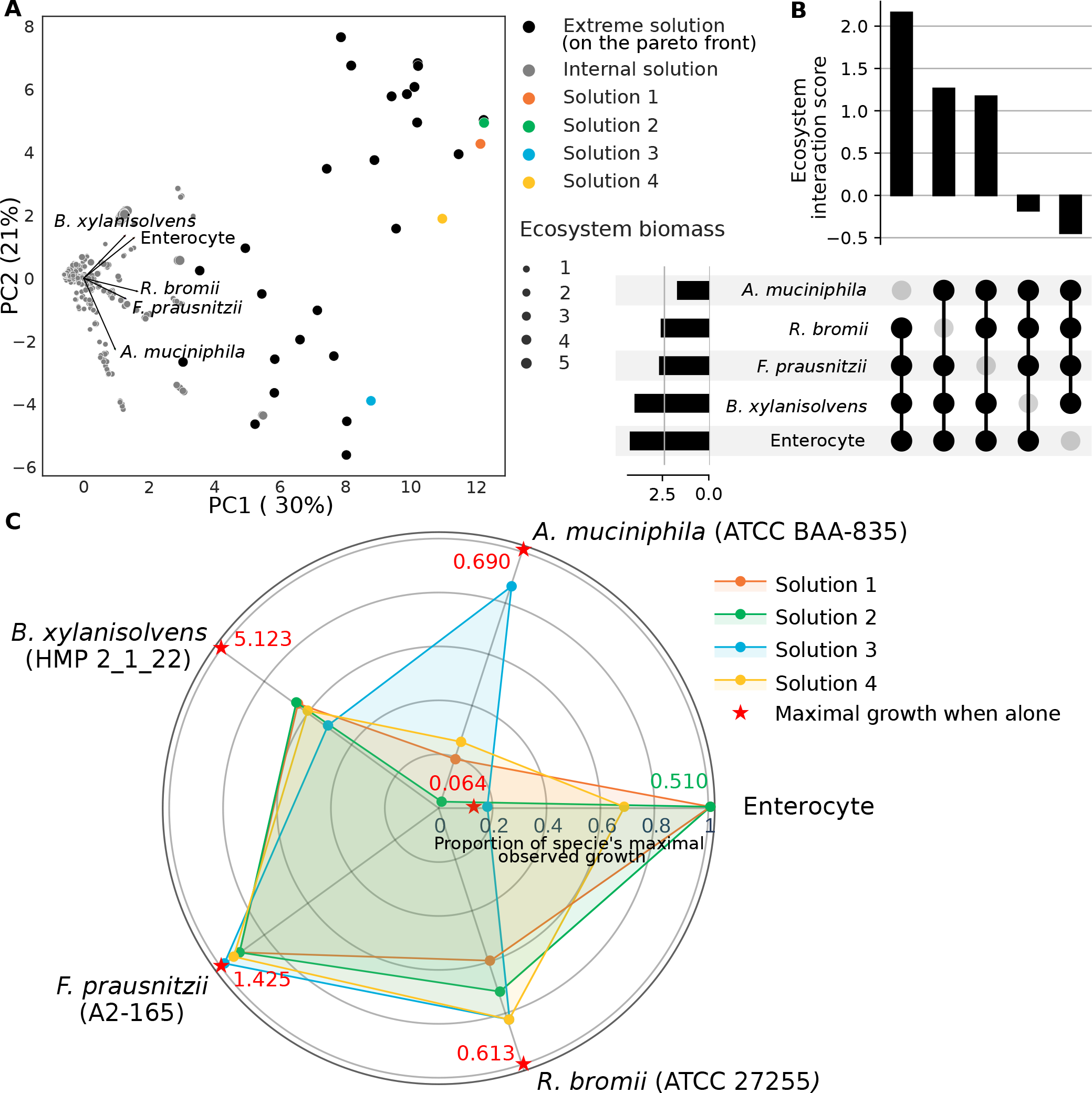
Multi-objective community metabolic modeling of five organisms in an ecosystem (enterocyte, *Akkermansia muciniphila, Bacteroides xylanisolvens, Faecalibacterium prausnitzii*, and *Ruminococcus bromii*). **(A)** Principal component analysis of solutions defined by the objective value of each organism. The point’s size is proportional to the sum of this solution’s objective values. Grey points result from a random sampling in the solution space of the ecosystem. Black and colored points are extreme solutions defining the Pareto front. Colored points are a selection of extreme solutions from the Pareto front where every organism (objective) has a non-null objective value, representing cohabiting metabolic phenotypes for the ecosystem where each organism can grow or maintain itself. **(B)** Upset diagram plot of the impact of the absence of each organism on the ecosystem interaction score. Scaling the interaction score from bi-objective to four-dimensional ecosystems is required to calculate hypervolumes instead of the area under the curve (compared to Figure 1) but follows the same principle (see Methods). **(C)** Visualization of the four cohabiting metabolic phenotypes for the ecosystem (solutions where every organism had a non-null objective value). The axes are objective values for the corresponding organism, normalized by its highest observed objective value (in the ecosystem or alone).

Inferring the interaction score for the ecosystem (See Methods) is informative of its overall interaction. Here, an ecosystem interaction score of 1.70 implies that the modeled organisms are dominated by mutualist interaction compared to competition. However, this lacks precisions about the role of each actor in the interaction. As a strategy to highlight the impact of a given organism on the ecosystem’s interaction potential, we calculated the score of smaller ecosystems after the removal of each organism. Therefore, we measured the score for five reduced ecosystems (removing a different organism each time) to better understand the role of each organism in the community’s interplay (Figure 4B). Removing the enterocyte or *B. xylanisolvens* from the ecosystem yielded negative scores (-0.44 and -0.18). This suggests that they are involved in a beneficial relationship, raising the overall score of the ecosystem. The score remained positive when removing *F. prausnitzii* or *R. bromii* (1.17 or 1.26). The highest interaction score was observed when *A. muciniphila* was absent from the ecosystem (2.16). Since one cannot mathematically compare these four-dimensional scores to a five-dimensional score when all organisms are present in the ecosystem, we cannot conclude whether *Akkermansia muciniphila* exhibits a weaker but still beneficial interaction or if it decreases the overall score of the ecosystem. Still, it is the bacteria with the least favorable effect on the score. The impact of each organism on the ecosystem score is coherent with their disposition on the PCA (Figure 4A), concluding that PC2 is correlated with the interaction score.

Four extreme solutions were selected where no organism had a null objective value (Figure 4C). The maintenance of the enterocyte was highly favored by the presence of bacteria, with an improved objective value in all selected solutions. However, it is noteworthy that the growth of the bacteria within the ecosystem is reduced compared to their maximum growth potential when in isolation. The objective values of the enterocyte and the *B. xylanisolvens* are correlated across solutions. Besides, *R. bromii, F. prausnitzii*, and particularly *A. muciniphila* seemed to follow an opposite trend. In this situation, *B. xylanisolvens* seems involved in a mechanism essential to the enterocyte’s improved maintenance. *B. xy-lanisolvens* is a complex polysaccharide degrader (32, 33), making carbohydrates and folate available to the enterocyte, which may explain this positive interaction. However, when reaching optimum, objectives of the enterocyte and *B. xylani-solvens* were antagonists to the growth of *A. muciniphila. A. muciniphila* is a next-generation probiotic associated with intestinal and systemic health, thought to preserve gut barrier function in the intestinal tract (34). The known metabolic action of *A. muciniphila* on the gut epithelium is through short-chain fatty acids (SCFAs) production (35). However, short-chain fatty acids’s impact is prevalent in the colon (36), and is not modeled in the enterocyte used in this study.

## Discussion

Deciphering the interactions between epithelial cells and gut microbes is a crucial step toward a better understanding of human health. This motivates their exploration through the use of GEMs to uncover potential metabolic interplays. This study uses multi-objective modeling to capture the trade-offs between the host’s and gut bacteria’s objectives, and predict putative metabolic interaction mechanisms. We summarized this complex information by a generic score that indicates overall collaborative, competitive, or neutral interactions between organisms, as well as the quantification of this potential interaction. Known probiotics (15) were associated with a higher interaction score with the enterocyte, raising interest in the score’s potential as a screening tool to assess interactions between organisms. Interaction types were computed from the Pareto front’s shape, further characterizing the interaction. The interaction score varied depending on the diet imposed on the ecosystem. Notably, the protein diet favored a more mutualist interaction, while the Western diet raised competition. However, the nutritional constraints applied to model the diets were under-constrained. Specifically, nutrients with known concentrations were adequately restricted, while unspecified nutrients were freely available in the lumen. Well-defined nutritional constraints are time-consuming to calibrate and are based on elaborated knowledge of the modeled medium (37). Designing a diet allowing all 331 bacteria models to grow with the enterocyte without biological reference is time-consuming and imprecise. Given that under-constrained ecosystems still highlight the influence of the diet on metabolic interactions, we anticipate that further research, with more targeted subjects and strictly defined diets, will yield predictions of higher accuracy. Overall, the present predictions are dependent on the models’ quality. We, therefore, expect that, as scientific effort lessens classic modeling limitations (See S4), the exactitude of the predictions will improve.

*Lactobacillus rhamnosus* GG (LGG), a known probiotic (26), has been identified *in silico* as such through a high interaction score and a mutualistic interaction type. This predicted interaction was explored to identify the metabolic mechanisms behind it. This study identified nutrients driving the beneficial interaction between LGG and the host, highlighting the importance of vitamins, amino acids, and oligo-elements, already recognized as central cross-feeding metabolites (7). Moreover, cross-feeding was predicted, implying the desulfatase of choline sulfate by LGG, making choline available for the enterocyte while using sulfate ions to grow. As shown before, choline metabolism into phosphatidylcholine participates in cell membrane synthesis and lipid metabolism (29). Choline can be converted to trimethylamine (TMA) by the colonic microflora, which, once absorbed by the colonocytes and transported to the liver, can be oxidized in trimethylamine N-oxide (TMAO) (38). TMAO is associated with various diseases, such as cardiovascular disease, when found in high concentrations (39). However, due to the use of overapproximations for nutritional constraints, the predicted mechanisms are also prone to be over- or under-estimated. In this context, as choline sulfate availability is modeled in excess, we anticipate that the exchange of choline would occur in smaller quantities than initially predicted, thereby preventing the accumulation of high concentrations of TMAO in the liver. Presently, *in vitro* experiments are essential to confirm this cross-feeding.

The score and analysis developed in this study are tools to explore ecosystems’ metabolic interplays. Their application to communities of interest, such as a consortium of gut bacteria associated with a health condition, could participate in identifying pathogenic or therapeutic pathways. Similarly, the health impact of dietary products with known bacterial composition, like cheese or yogurt (40), could be further explored. Moreover, assessing which type of epithelial cell interacts the most with a species could enrich our knowledge of how different gut locations build different communities (41). As an illustration, we expect *Akkermansia muciniphila* to reach a higher interaction score with a colonocyte than the small intestine epithelial cell used in this work. While this framework was demonstrated here on metabolic interactions between the human gut epithelium and the microbiota, its application is generic enough to be relevant for various ecosystems and contexts. Tumors are accompanied by bacteria, which promote or suppress cancer based on the situation (42). Applied in this tumor-microenvironment ecosystem, a positive score would highlight pathogenic interaction, opening the way for adapted treatments, like targeted antibiotics. Finally, this can be applied to non-health-related fields, such as the study of ocean and soil ecosystems. Specifically, it can allow the study of uncultured strains through the modeling of Metagenome-Assembled Genomes (MAGs).

In this work, we converge with previous results of Heinken and Thiele (24), using multi-objective optimization to provide a mathematical proof of symbiosis. Indeed, we demonstrate here that two organisms in nutritional co-limitation can attain better fitness than alone. This observation justifies the development of tight multi-species communities in various ecological contexts, as their cohabitation leads to better survival. Additionally, we show that the diet modulates the importance of this mutualism. Compared to an open environment such as the ocean, the gut is an enclosed habitat where diet is the principal nutritional input. On the other hand, the microbiota regulates food intake (in terms of quantity and quality) through the gut-brain axis (43). Therefore, our nutrition forges our microbiota’s interactions and fitness, shaping its composition (44), while the microbiota influences our nutrition (43). Controlling the microbiota’s early assembly and ecology through an adapted diet is key to favoring a self-maintaining healthy composition (45). This encourages pursuing research on the gut ecosystem metabolism and how microbiota and nutrition interact and impact human health.

## Methods

### Genome-Scale Metabolic Models

The **331 bacteria models** used in this study are CarveMe reconstructions extracted from the publicly available EMBL GEMs database (16). These bacteria were selected based on their description as gut bacteria in the Virtual Metabolic Human (VMH) database (14). The bacteria described in the VMH database’s metadata as “Pathogen,” “Opportunistic pathogen,” or “Putative Pathogen” were joined in the “Pathogen” ecological type (n = 46). The ones described as “Probiotic” or “Probiotic potential” were joined in the “Probiotic” ecological type (n = 18). The “Commensal” (n = 123) and “uncharacterized” (n = 144) ecological types were conserved as is.

The **small intestine epithelial cell (sIEC)** model relies on previously published results (46) and includes 1282 reactions and 844 metabolites. The namespace of the exchange reactions (i.e., the nomenclature of the model elements) was adapted to the EMBL models (i.e., BiGG’s namespace) for model compatibility. The sIEC model includes two external compartments: the blood and the lumen. In constructing the ecosystem model, the reactions controlling the apparition and disappearance of metabolites in the blood were not considered exchange reactions, in contrast with those in the lumen.

### Ecosystem model: pool compartment, diet

bacterial model, and the sIEC’s model were joined to exchange metabolites through a **pool compartment** to build an ecosystem model. Mathematically, all models’ stoichiometric matrices were diagonally assembled into a new one, in which a pool compartment was added. The original exchange reactions of the models became transport reactions (TR) from the organism’s external compartments to the pool compartment. All exchange reactions in at least one of the models were duplicated to form the pool’s exchange reactions (ER) group. TR were unconstrained to enable free transit of the metabolites between the pool and the organism’s external compartment, and the media constraint was applied by restricting the bounds of ER.

This metabolic model of the ecosystem was built using a Python version of mocba (23) (i.e. mocbapy), and further constrained to fit the nutritional conditions described above on the pool compartment.

In this study, the unconstrained diet consists of the absence of constraint on the exchange reactions of the pool. The Western diet (WD) and Protein diet (PD) were extracted from Heinken et al. (24). WD is high in simple sugar (47%) and fat (35%) and is low in fiber (3%) and proteins (15%). PD is high in protein (30%) and balanced in fat (20%), simple sugar (25%), and fiber (25%). For WD and PD conditions, exchange reactions between the sIEC and the blood were constrained based on the Average American Diet (AAD) (46).

When constraining the pool’s content strictly to the diet information, with every other reaction blocked, the bacteria could not grow. Therefore, the exchange reactions not described in the diet were left unconstrained.

### Multi-objective linear programming: Pareto front

Multi-objective linear programming is a mathematical optimization technique that optimizes multiple conflicting objectives by finding optimal solutions representing the trade-offs between the different objectives. This trade-off is formally a Pareto front describing the optimal behaviors of the resulting metabolic ecosystem, in the sense that no increase in an objective can be done without affecting (i.e., decreasing) others. For identifying the Pareto front for an ecosystem composed of multiple metabolic models, we used the mocbapy python package. It translates the metabolic model of the ecosystem into a multi-objective linear problem and solves it using a Python adaptation of Bensolve (47, 48) to identify the Pareto front. This Pareto front is a set of extreme points in the objective space. This case study’s objectives are the bacteria and the sIEC biomasses.

### Linear problem formalisms

A metabolic model comprises a stoichiometric matrix (**S**) representing metabolite-reaction relationships, fluxes (**v**) representing reaction rates, and bounds (*l*_*i*_ and *u*_*i*_) defining flux constraints. A solution for a usual mono-objective FBA is obtained by solving the following linear problem:

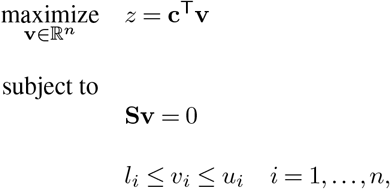

where **c**^T^**v** is a linear combination of fluxes representing the objective function.

In a multi-objective linear problem, the stoichiometric matrix is organized in several compartments, being able to exchange metabolites through a pool compartment. There are as many optimized objective functions as organisms modeled in the ecosystem. As described in Budinich et al. (23), the multi-objective linear problem solved in this instance can be defined as

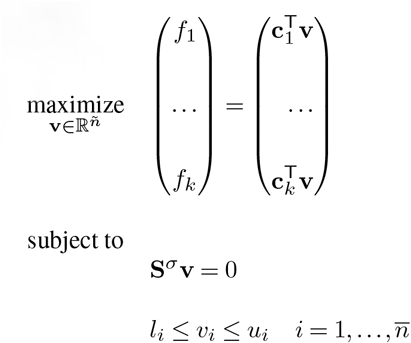

where (*f*_1_, …, *f*_*k*_)^T^ are the objective functions of the *k* organisms and *n* is the total number of reactions (*i*.*e*., the sum of reactions of each organism and exchange reactions from the pool compartment).

### Interaction score

The Pareto front of **bi-objective** problems is described in a two-dimensional space, each axis describing the possible values for one objective. To calculate an interaction score, the values of each dimension are normalized by the maximal value of their respective objective when the organism is alone. The maximal growth of the bacterium and the enterocyte alone are added to the Pareto front as (0, 1) and (1, 0), respectively. The interaction score is inferred from the area under the normalized Pareto front (*AUC*_*P*_) curve, from which we subtract the area under the curve of the non-interaction front (*AUC*_*NI*_).

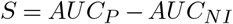

In a problem of **more than two objectives**, the same normalization is applied to the extreme points defining the Pareto front. The points of maximal growth when alone of each organism are added accordingly, as well as the origin point (a vector with zero values for all dimensions). The convex envelope of this set of points and its hypervolume are calculated using Scipy’s ConvexHull function (49). The interaction score is the hypervolume, thus calculated from which is subtracted the non-interaction hypervolume.

When alone, all models were solved with CPLEX. Multi-objective problems were solved with benpy, a Python adaptation of bensolve.

### Interaction types

In two dimensions, the various forms taken by the Pareto front were discriminated into categories based on four conditions. Sign: Sign of the score. E+: A solution exists on the Pareto front where the biomass value of the enterocyte in the ecosystem is higher than that of the enterocyte alone. B+: A solution exists on the Pareto front where the biomass value of the bacterium in the ecosystem is higher than the biomass value alone. E+B+: A solution on the Pareto exists where the biomass values of both organisms are at their highest. They were not studied in problems of higher dimensions.

The various forms taken by the Pareto front were discriminated into categories based on four conditions.

1. **Sign**: Sign of the score.
2. **E+**: A solution exists on the Pareto front where the biomass value of the enterocyte in the ecosystem is higher than that of the enterocyte alone.
3. **B+**: A solution exists on the Pareto front where the biomass value of the bacterium in the ecosystem is higher than the biomass value alone.
4. **E+B+**: A solution on the Pareto exists where the biomass values of both organisms are at their highest.

**Table 1.** Interaction types inferred from Pareto appearance)

| Sign | E+ | B+ | E+B+ | Interaction type |
| --- | --- | --- | --- | --- |
| + | False | False | False | Neutralism |
| - | False | False | False | Competition |
| + | True | False | False | Favors enterocyte |
| + | False | True | False | Favors bacteria |
| + | True | True | False | Limited mutualism |
| + | True | True | True | Mutualism |

### Sampling of the Pareto front

A homogeneous sampling of 100,000 solutions was performed along the Pareto front, describing the interaction between LGG and the enterocyte. Each sample was obtained by conducting a Flux Balance Analysis (FBA) with fixed biomasses of both organisms at the chosen Pareto point. The ecosystem biomass (sum of each biomass) was added to the resulting sampling, and a Spearman correlation matrix was generated. The exchange reactions with a correlation higher than 0.95 (secreted) or lower than -0.95 (uptaken) with the ecosystem biomass were considered drivers of the mutualistic interaction between LGG and the enterocyte (Fig 3B).

### Exchanged metabolites and obligatory reaction

Exchanged metabolites are metabolites for which the secretion by an organism is correlated (Spearman, corr > 0.5) to their absorption by the other organism. Obligatory reactions cannot be inactive based on a Flux Variability Analysis (FVA), which determines the upper and lower possible value for flux in a given condition.

### Building a five-dimension ecosystem

Based on Shetty et al. (31), 16 bacteria strains were selected to represent a minimal human gut microbiome. First, the sequenced genome of the strains were annotated with prokka (50). Then, CarveMe (16) was used to generate metabolic models for each organism (See genome references in Table S3). For computational reasons, a Multi-objective problem could be solved with up to 10 bacteria models in addition to the enterocyte (Figure S3). We observed that some bacteria were in total competition, meaning that if one bacteria was growing, the others had null growth. This led to the selection of four cohabiting bacteria: *Akkermansia muciniphila* ATCC BAA-835, *Bacteroides xylanisolvens* HMP 2_1_22, *Faecalibacterium prausnitzii* A2-165 and *Ruminococcus bromii* ATCC 27255. They were joined with the enterocyte in an ecosystem constrained with a WD, and the extreme solution points in the inferred Pareto were retrieved.

### Five-dimensional principal component analysis (PCA)

In the five-dimensional ecosystem, 34 extreme points were identified on the Pareto front using mocbapy. To integrate solutions embedded in the solution space, the model was converted from mocbapy format to cobrapy. Then, the sampling function from cobrapy was applied to retrieve 3000 random solutions. In each solution, only the objective values for each organism were kept for the PCA. The data was processed and the model was built using sckitlearn (51) tools (preprocessing.StandardScaler() and decomposition.PCA()). The explained variance of the five principal components were 30%, 21%, 18%, 17% and 14%, respectively. As the principal components three and four had non-negligeable variance explanations, the knee point was found using the kneed python package (52). The knee point is the point of maximum curvature in a function. Only the first two principal components were explored as the knee-point was equal to two.

## Quantification and Statistical Analysis

All statistical tests were realized using Python 3.7.13 (53). Significance was determined from a p-value inferior to 0.05. The comparison of interaction score values for ecological types (Figure 2C) was assessed by pairs of ecological types with a Mann-Whitney U test using numpy.stats.mannwhitneyu(). The ecological types comprised n = 46 pathogens, n = 18 probiotics, n = 123 commensals and n = 144 uncharacterized. Every correlation was calculated with pandas.corr(method = “spearman”). A correlation was considered of interest if its absolute value was equal or over 0.95. To calculate the score, the AUC (in two-dimensional Pareto fronts) was calculated with scikit learn 1.0.2 (51) (sklearn.metrics.auc) and the hypervolume (Pareto fronts of three or more dimensions) was calculated with scipy 1.7.3 (49) (scipy.spatial.ConvexHull.volume). The figures were built using matplotlib 3.5.2 (54), seaborn 0.11.2 (55) and plotly 5.10.0 (56).

## Resource availability

### Lead contact

Further information and requests for resources should be directed to and will be fulfilled by the Lead Contacts, Anna Lambert or Damien Eveillard.

## Materials availability

This study did not generate new materials.

## Data and Code Availability

The 331 gut bacteria models were CarveMe reconstruction from the EMBL GEMs database available at: EMBL_GEMs

The small intestine intestinal cell model is available at: In silico reconstructions - Thiele lab

The minimal microbiome models built for this paper are available at: github - MO_GEMs_Score

All original code has been deposited at github - MO_GEMs_Score and is publicly available as of the publication date.

## ACKNOWLEDGEMENTS

AL benefits a doctoral fellowship managed by Bba Milk Valley, dairy industrial association. We thank Regional Councils of Bretagne (grant no. 19008213) and Pays de la Loire (grant no. 2019-013227) for their financial support through the interregional project PROLIFIC. The bioinformatics core facility of Nantes (BiRD, Biogenouest), University of Nantes, France, provided computational support.

## Author contributions

Conceptualization, A.L., D.E., S.C. and M.M.; Methodology, A.L., D.E. and S.C.; Software, A.L. and M.B.; Formal Analysis, A.L., Investigation, A.L.; Writing - Original Draft, A.L.; Writing - Review & Editing, A.L., D.E., S.C, M.M. and M.B.; Visualization, A.L.; Supervision, D.E., S.C. and M.M.

