## Supplementary figures and images for "Community metabolic modeling of host-microbiota interactions through multi-objective optimization"

### Figure S1

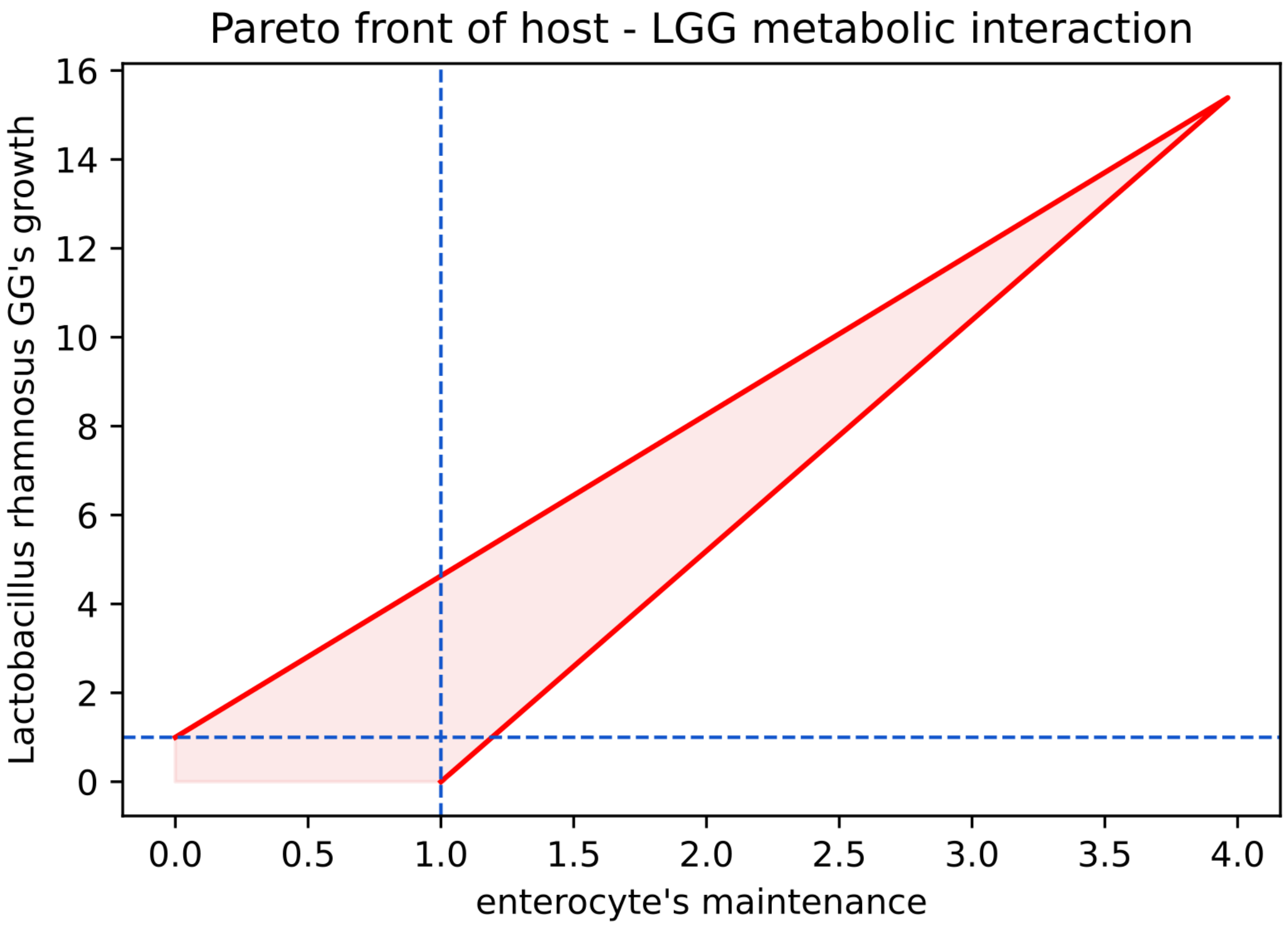

### Figure S2

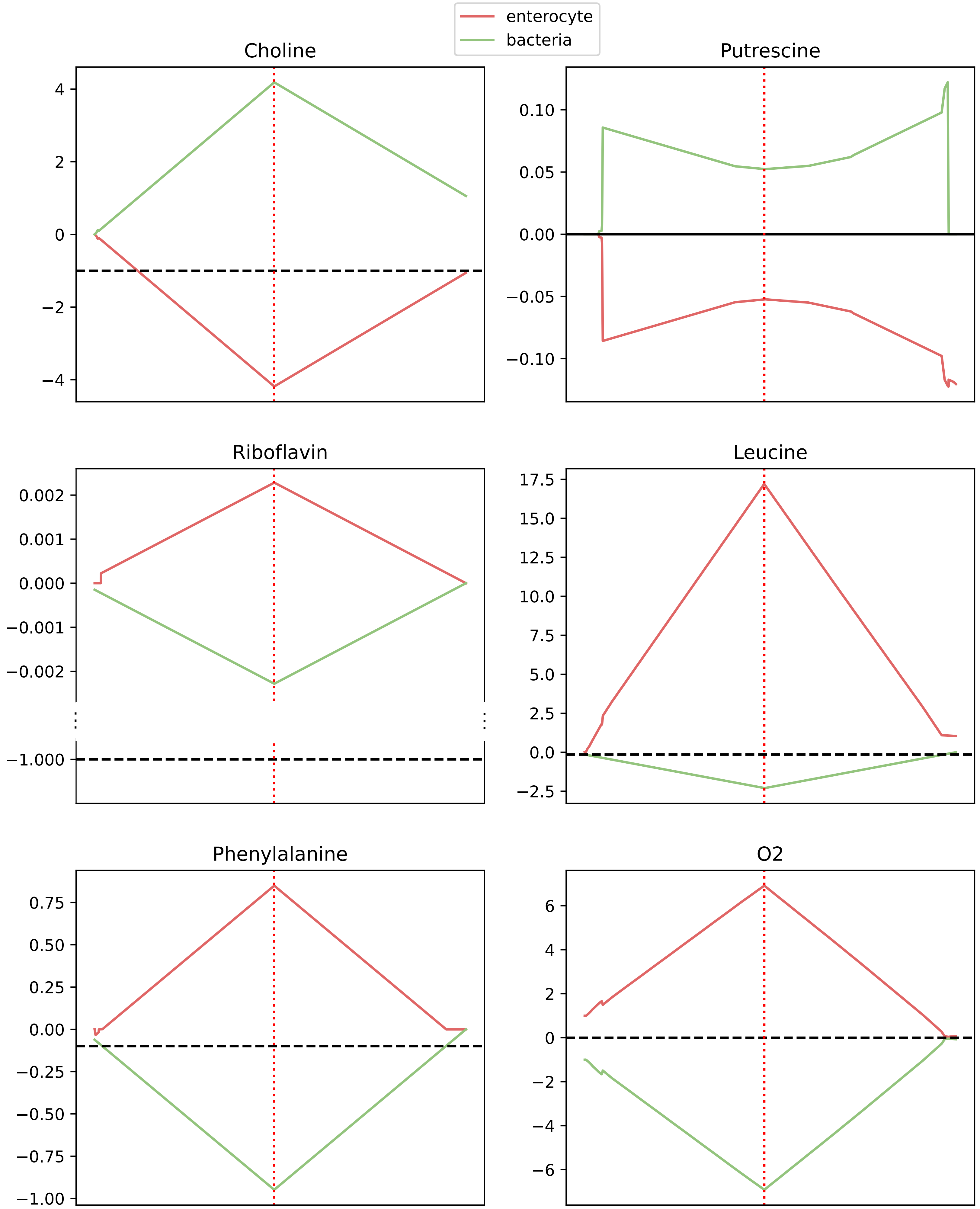
