## Supplementary material for "Community metabolic modeling of host-microbiota interactions through multi-objective optimization": Figure S3

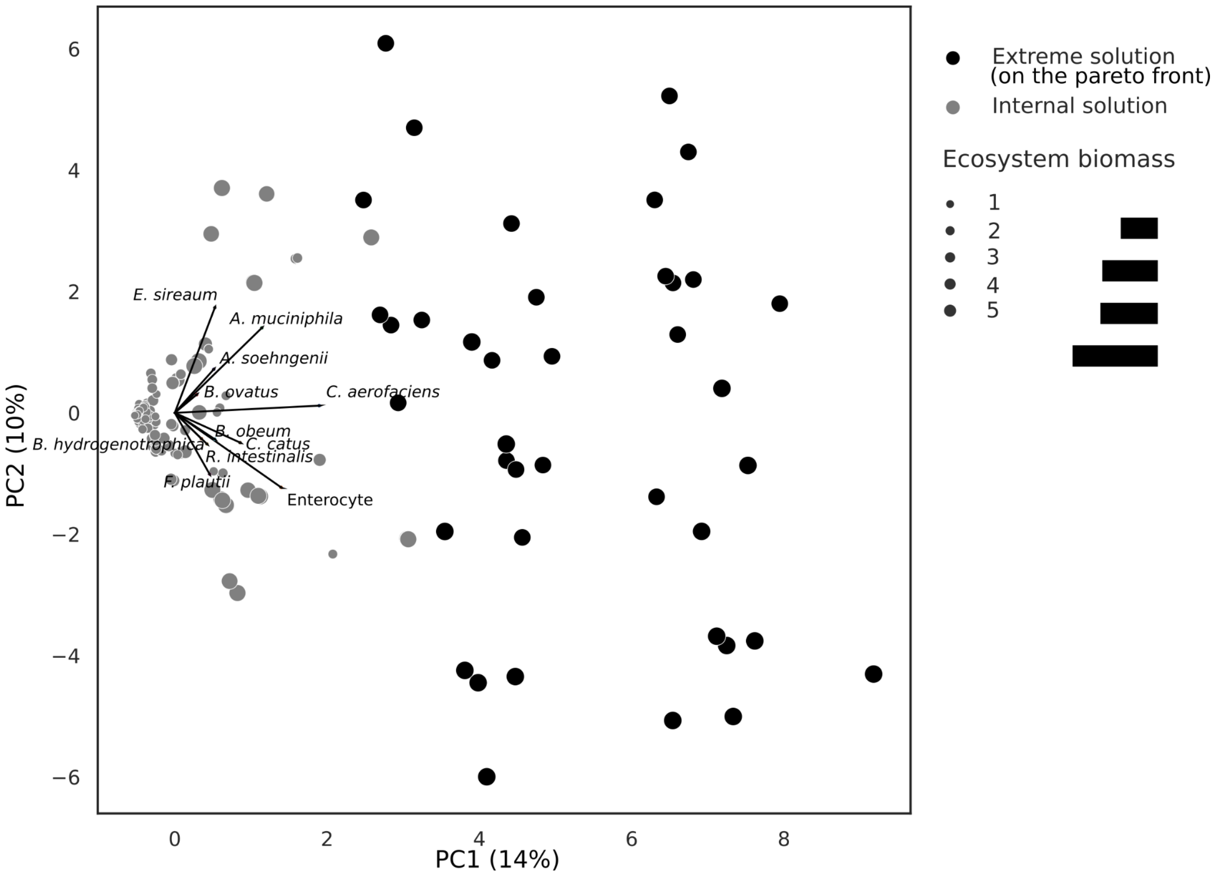


**Figure. S5 | PCA on the objective space of the multi-objective community metabolic modeling of 11 organisms (enterocyte, *A. muciniphila, A. soehngenii, B. hydrogenotrophica, B. obeum, B. vatus, C. aerofaciens, C. catus, E. sireaum, F. plautii, R. intestinalis*) in an ecosystem.**

Computational resources enabled the multi-objective analysis of 10 gut bacteria joined with the enterocyte. Among the 16 strains representing a minimal gut microbiome according to Shetty et. al. (1), we selected the ten strains from with the smallest niche overlap (*Akkermansia muciniphila* MucT/ATCC BAA-835, *Anaerobutyricum soehngenii* L2-7/DSM 17630, *Blautia hydrogenotrophica* DSM 10507, *Blautia obeum* DSM 25238, *Bacteroides ovatus* HMP strain 3_8_47FAA, *Collinsella aerofaciens* DSM 3979, *Coprococcus catus* ATCC 27761, *Eubacterium sireaum* DSM 15702, *Flavonifractor plautii* HMP strain 7_1_58FAA, *Roseburia intestinalis* DSM 14610).

To explore how organisms influenced each other's objectives, we realized a PCA on both the extreme points of the Pareto front and 3000 random solutions sampled in the objective space (Figure S5). Similarly to the results observed in the five-dimensional ecosystem described in the core paper, PC1 is overall associated with the ecosystem biomass' value. Here again, each organism participates in the increase of biomass production in the ecosystem. However, the variance explained by PC1 and PC2 only reach 24% and the high-dimensional nature of the data makes it difficult to raise many conclusions from the directions taken from each organism on the PCA. Additionally, every solution on the Pareto front implied the absence of growth for at least one organism of the ecosystem, implying total competition between some organisms. Finally, the interaction score was calculated for a reduced ecosystem, removing one organism each time, to evaluate the impact of each organism on the overall positive or negative interaction of the ecosystem. This resulted in only scores of -1, hinting towards a highly competitive ecosystem.

1. Shetty SA, Kostopoulos I, Geerlings SY, Smidt H, de Vos WM, Belzer C. Dynamic metabolic interactions and trophic roles of human gut microbes identified using a minimal microbiome exhibiting ecological properties. ISME J. sept 2022;16(9):2144‑59.
