## Supplementary material for "Community metabolic modeling of host-microbiota interactions through multi-objective optimization": S4

### Limitations

As previously stated in the discussion with nutritional constraints, models are approximative representations, and present limitations. The models used in this study were automatically reconstructed from a genome using the CarveMe pipeline (1). Due to their dependence on genome completion and lack of curation based on the literature on the specific strain, these models are more susceptible to diverging from chemical knowledge, potentially leading to the erroneous representation of metabolic pathways.

Here, the cross-feeding predicted between LGG and the enterocyte relies on the ability of LGG to metabolize choline sulfate into choline and sulfate. Choline sulfatase is a protein synthesized from the gene BetC. This gene was found in many bacteria but not in LGG (2). This is a reaction added through gap filling, meaning that it was necessary for the growth of LGG *in silico* although not found in its genome. From this information, we can hypothesize that LGG is able to perform an analogous reaction utilizing an alternate enzyme. Another possibility is that another pathway is necessary for LGG to grow, but a lack of information resulted in the addition of the choline desulfatase pathway to compensate for its absence. Overall, this reveals a gap in knowledge in the metabolism of LGG, requiring further investigation.

Additionally, the prediction of this method highly depends on the defined objectives. Indeed, the model would benefit from including more than maintenance in the enterocyte's objective function, as it is a differentiated cell with a specific role for the human body. Consequently, with the ongoing evolution of objective definitions and comprehensive curation, a corresponding improvement is expected in the score's accuracy.


1. Machado D, Andrejev S, Tramontano M, Patil KR. Fast automated reconstruction of genome-scale metabolic models for microbial species and communities. Nucleic Acids Res. 6 sept 2018;46(15):7542‑53.
2. Cregut M, Durand MJ, Thouand G. The Diversity and Functions of Choline Sulphatases in Microorganisms. Microb Ecol. 1 févr 2014;67(2):350‑7.
